# Detection of Japanese Encephalitis (JE) Virus in Piggery effluent from an Australian Farm Prior to Outbreaks of JE

**DOI:** 10.1101/2025.09.22.677956

**Authors:** Warish Ahmed, Metasebia Gebrewold, Leah G. Starick, Regina Fogarty, Kirsty Richards, David Williams, Stuart L. Simpson

## Abstract

Japanese encephalitis virus (JEV), a mosquito-borne orthoflavivirus, raises concerns about its seasonal re-emergence. Pigs are a major amplifying host and JEV infection can manifest as significant reproductive disease and losses, necessitating robust surveillance. This study evaluated effluent surveillance for early JEV detection in piggery effluent from a Victorian farm between December 2024 and March 2025. Effluent samples were tested using JEV-specific real-time reverse transcription (RT)-PCR, with positive detections found on four separate sampling days, despite an absence of clinical signs in livestock through the testing period. Subsequent veterinary investigations for JEV in litters born on 17/05/2025 (suspect cases only) aligned with a positive effluent sample collected during the estimated JEV infection exposure periods of the sows. With the single clinically confirmed JEV case (which farrowed on 1/06/2025) collection of the same positive effluent sample fell just outside, but near, the estimated exposure period for the affected sow. These findings highlight ability of effluent monitoring to detect JEV infection 2 to 4 months before clinical manifestations present, offering potential for a non-invasive, herd-level early warning system. Intermittent detections may suggest limitations in grab sampling and low viral loads in effluent samples. Integrating effluent surveillance with veterinary clinical testing of litters suspected of being exposed to JEV or during high-risk periods could enhance JEV management in Australia’s enzootic regions, supporting One Health strategies.

## Introduction

Japanese encephalitis virus (JEV), a mosquito-borne virus of the genus *Orthoflavivirus* (family *Flaviviridae*), remains a leading cause of viral encephalitis across Asia and the western Pacific (Postler et al. 2023; Mackenzie and Smith 2024; Yun et al. 2025). Since its emergence in southeastern Australia in 2022, concerns have grown regarding its potential for seasonal re-emergence, particularly under favourable climatic conditions. JEV is sustained in an enzootic cycle involving water birds and mosquitoes, particularly those belonging to the *Culex* genus, with pigs and wild birds acting as amplifying hosts (Mulvey et al. 2021). Among domestic animals, pigs are significant due to their role in viral amplification and their susceptibility to reproductive disorders disease following infection (McLean and Graham 2022).

Current JEV control strategies comprising human vaccination, mosquito management, and public education may be inadequate in areas with high mosquito activity, limited vaccine uptake, or where swine vaccination is not implemented (Park and Lee 2025). Notably, whilst vaccine development work is in progress, no JEV vaccine is currently approved for use in pigs in Australia (Harrison et al. 2024). Consequently, enhanced surveillance systems are essential for early detection and timely response, particularly in regions experiencing new or recurrent JEV activity (Sharan et al. 2023). Wastewater surveillance has emerged as a valuable population-level monitoring tool for infectious diseases in humans such as SARS-CoV-2 and influenza A virus (IAV) (Viviani et al. 2025).

A recent study demonstrated the potential of effluent and environmental water surveillance for detecting JEV in piggery settings (Ahmed et al. 2025). That study emphasized the importance of regular monitoring, especially during summer months when specific environmental and climatic conditions favour mosquito-borne virus transmission. This approach enables early identification of viral shedding prior to the appearance of clinical signs in farrowed sows, as infection during gestation typically results in *in utero* death of fetuses that is not detected until farrowing (piglet birth) - an outcome that may lag behind transmission and viraemia by weeks.

In this study, we report the early detection of JEV in piggery effluent samples collected from a farm at a time when none of the livestock exhibited signs of infection. This was subsequently followed by a confirmed JEV outbreak. These findings underscore the utility of effluent-based surveillance as an early warning system for JEV activity in high-risk agricultural environments. As global health systems increasingly adopt integrated One Health approaches, environmental surveillance tools such as effluent monitoring offer a practical and proactive strategy for mitigating zoonotic disease risks at the human–animal–environment interface. Here, we evaluate effluent surveillance as a complementary tool to detect JEV prior to clinical outbreaks.

## MATERIALS AND METHODS

### Effluent sampling and concentration of viruses

Effluent samples were collected from a pig farm in Victoria between 17/12/2024 and 11/03/2025. Eight effluent grab samples (50 to 200 mL) were collected on eight different days from a pit receiving effluent from flush channels that exited pig sheds. Pits are in-ground tanks storing at least one day of effluent from multiple sheds.

### Nucleic acid concentration and extraction from effluent

A 50 mL effluent sample was centrifuged at 4,750 *g* for 20 min, and the remaining sample was archived at -20°C (Ahmed et al. 2025). The pellet resulting from the centrifugation step was collected from the 50-mL tube and transferred into a 7-mL bead-beating tube for nucleic acid extraction. Nucleic acid extraction from the pellet (effluent solid fraction) was performed using the RNeasy PowerWater Kit (Cat. No. 14700–50-NF, Qiagen) according to the process described elsewhere (Ahmed et al. 2025). Nucleic acid was eluted in 150 μL of DNase- and RNase-free water and stored at −20°C for 72 to 96 h prior to RT-PCR analysis.

### RT-PCR assays

The detection of JEV was carried out using RT-PCR assays targeting the JEV non-structural protein--1 (Universal JEV assay) and membrane (JEV G4 assay) genes of JEV (Shao et al. 2018; Wang 2022, pers. comm.), as previously described (Ahmed et al., 2025). The Universal JEV assay enables the detection of all JEV genotypes, while the JEV-G4 assay specifically detects JEV genotype 4, the current genotype circulating in Australia (Mackenzie et al. 2022; Ahmed et al. 2025). Initially, Universal JEV RT-PCR was conducted, but it was later replaced with the JEV G4 RT-PCR assay due to the latter’s increased sensitivity (Freddi et al. 2025, under review).

RT-PCR amplifications were performed with a 20 µL reaction mixture containing 5 µL of TaqMan™ Fast Virus 1-Step Master Mix (Applied Biosystems, Thermo Fisher Scientific, MA, USA). A 5 μL nucleic acid sample was used in each reaction. Each run included triplicate nucleic acid samples, positive controls, and no-template controls. The RT-PCR assays were performed on a Bio-Rad CFX96 thermal cycler (Bio-Rad Laboratories, Hercules, California, USA).

### Oral fluid sampling and diagnostic testing of suspect litters

As an early season surveillance technique during the 2024-25 mosquito season, oral fluid samples were collected from pigs using chew ropes on a select number of piggeries at high risk of JEV infection (determined geographically and according to previous clinical history). On this farm the surveillance plan was to collect samples from up to 6 pens of young breeder pigs each week over the summer season.

Oral fluid collection via chew ropes is non-invasive and leverages a pig’s natural curiosity and chewing behaviour. Unbleached cotton ropes (three strand, 16 mm diameter, 50 cm length, Sydney Rope Supplies) were affixed to a fence or post at pig shoulder height using a cable tie and suspended in pens for approximately 15-20 min. During this time, pigs manipulated and chewed on the rope, depositing oral secretions. After removal from the pen, fluids were extracted by squeezing and wringing the rope in a plastic bag, allowing the fluid to pool at the base of the bag and dispensing the fluids into a sample tube. These samples were sent to the Victorian state veterinary laboratory, AgriBio for the detection of genetic fragment using RT-PCR and Flavivirus Antibody Enzyme-Linked Immunosorbent Assay (ELISA) analysis to detect antibodies against JEV, indicating prior exposure or immune response. Clinical samples from suspect litters i.e., foetuses, brain tissue samples and or placental material were periodically submitted directly to AgriBio by the herd veterinarian. Depending on the type of sample received, the on-duty veterinary pathologists performed additional processing as required. In cases where whole foetuses were submitted, brain tissue was extracted for further examination. All samples from suspect litters underwent RT-PCR testing for JEV.

No foetal material was available from suspect litters that farrowed on the 17/05/2025 but bloods samples were obtained from 3 sows for Flavivirus Antibody ELISA testing. Blood samples were collected by the herd veterinarian via venepuncture of the external jugular vein.

### Quality assurance/controls

A SARS-CoV-2 RT-PCR assay was applied to determine inhibition in extracted nucleic acid samples from effluents by seeding known copy number (10^4^) of gamma-irradiated SARS-CoV-2. Samples were considered to have no inhibition when the Cq values of effluent nucleic acid samples were within 2 Cq values of the reference Cq value (Staley et al. 2012). Nucleic acid extraction and RT-PCR setup were performed in separate laboratories to minimize potential sample cross contamination.

### Data interpretation

Samples were classified as positive for JEV by RT-PCR if amplification was detected in at least one out of three replicates within 40 cycles. Samples were considered indeterminate when the Cq value was between 40 to 45. A sample was considered negative when none of the three replicates showed amplification within 45 cycles.

## RESULTS

All RT-PCR negative controls consistently yielded negative results, verifying that no sample-to-sample contamination occurred in the RT-PCR. Positive controls consistently yielded positive amplification, validating the RT-PCR assay’s reliability. For the RT-PCR inhibition assay, the samples’ Cq values remained within the acceptable ±2 Cq range, confirming the absence of inhibition in the samples. Table 1 presents the results of JEV testing on the study farm over the period from 09/11/2024 to 03/06/2025, including effluent, oral fluid, and tissue samples using RT-PCR. In addition, oral fluid and blood/serum samples were also tested using ELISA. Effluent samples tested positive for JEV RNA on 14/01/2025, 21/01/2025, 18/02/2025, and 11/03/2025, while negative results were recorded on 17/12/2024, 04/02/2025, 11/02/2025, and 25/02/2025. None of the oral fluid samples tested over the period from 9/11/2024 to 11/03/2025 were found to be positive.

**Table 1:**
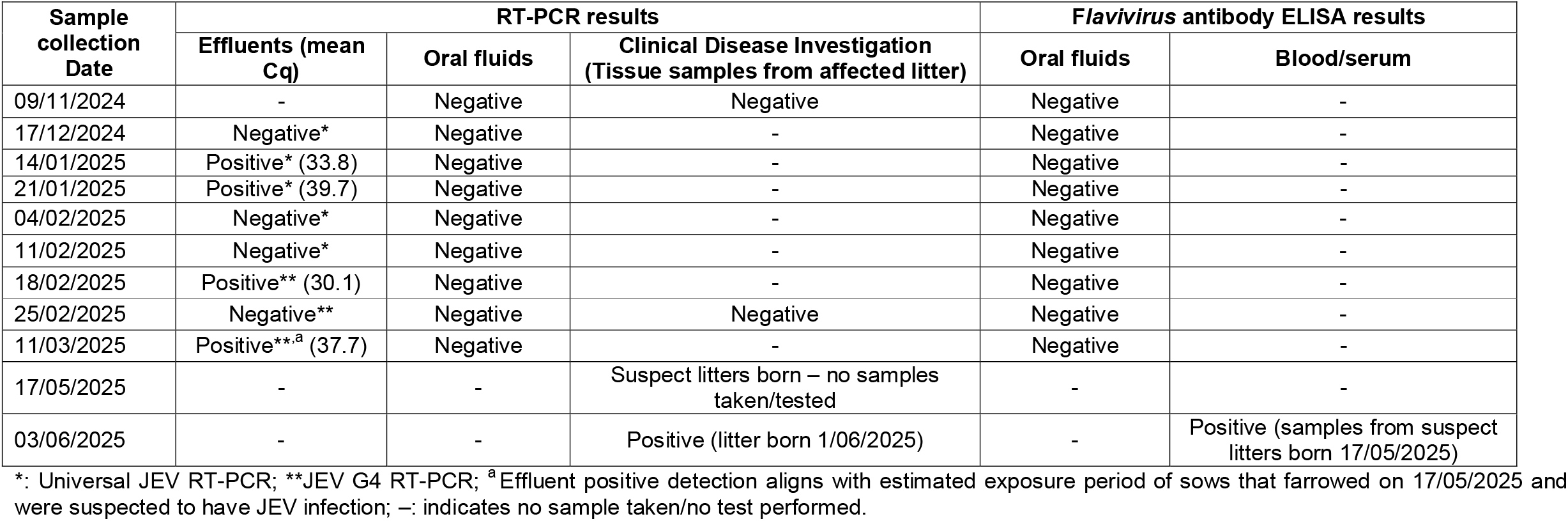
Detection of JEV in piggery effluent and oral fluid samples using RT-PCR and ELISA.

| Sample collection Date | RT-PCR results |  |  | Flavivirus antibody ELISA results |  |
| --- | --- | --- | --- | --- | --- |
|  | Effluents (mean Cq) | Oral fluids | Clinical Disease Investigation (Tissue samples from affected litter) | Oral fluids | Blood/serum |
| 09/11/2024 | - | Negative | Negative | Negative | - |
| 17/12/2024 | Negative* | Negative | - | Negative | - |
| 14/01/2025 | Positive* (33.8) | Negative | - | Negative | - |
| 21/01/2025 | Positive* (39.7) | Negative | - | Negative | - |
| 04/02/2025 | Negative* | Negative | - | Negative | - |
| 11/02/2025 | Negative* | Negative | - | Negative | - |
| 18/02/2025 | Positive** (30.1) | Negative | - | Negative | - |
| 25/02/2025 | Negative** | Negative | Negative | Negative | - |
| 11/03/2025 | Positive**<sup>a</sup> (37.7) | Negative | - | Negative | - |
| 17/05/2025 | - | - | Suspect litters born – no samples taken/tested | - | - |
| 03/06/2025 | - | - | Positive (litter born 1/06/2025) | - | Positive (samples from suspect litters born 17/05/2025) |

\*: Universal JEV RT-PCR; \*\*JEV G4 RT-PCR; <sup>a</sup> Effluent positive detection aligns with estimated exposure period of sows that farrowed on 17/05/2025 and were suspected to have JEV infection; –: indicates no sample taken/no test performed.

No clinical cases were confirmed in the herd until 03/06/2025. The suspect litters born on 9/11/2024 and 25/02/25 were RT-PCR negative at the AgriBio laboratory. Blood/serum samples taken from 3 suspect sows that farrowed on the 17/05/2025 were Flavivirus Antibody ELISA positive which suggests previous infection that may relate to the clinical disease seen in those litters. Piglet tissue samples collected from a single sow with a suspect litter (farrowed 1/06/2025) were RT-PCR positive to JEV meeting the national definition of a confirmed case (Animal Health Australia 24/12/2024).

Reproduction data for the suspect litters indicated that sows were mated on 19/01/2025 and farrowed on 17/05/2025, with potential JEV infection exposure period from 23/02/2025 to 30/03/2025. This timing aligned with the 11/03/2025 RT-PCR positive effluent sample as per Table 2. The litter confirmed positive by RT-PCR on 03/06/2025 (sow mated on 06/02/2025, potential JEV infection exposure period from 13/03/2025 to 17/04/2025, farrowed on 01/06/2025 with PCR positive litter) had no corresponding effluent samples collected during the estimated at-risk period. The estimated at-risk period was inferred from the clinical presentation of affected piglets (e.g., foetal size and type including mummified, stillborn, or malformed).

**Table 2:** Estimated JEV exposure period and farrowing dates of suspect and confirmed litters in relation to effluent surveillance.

| Description | Mating date | Estimated exposure period | Farrowing (birth) date | Notes |
| --- | --- | --- | --- | --- |
| Suspect litters in May | 19/01/2025 | 23/02/2025 to 30/03/2025 | 17/05/2025 | JEV positive effluent samples collected on 11/03/2025 aligns with exposure period |
| Confirmed positive litter in June | 06/02/2025 | 13/03/2025 to 17/04/2025 | 01/06/2025 | Effluent sample collection did not continue through the potential at-risk period for this litter |

## DISCUSSION

The intermittent detection of JEV RNA in piggery effluent samples from 14/01/2025 to 11/03/2025, preceded confirmed veterinary diagnostic cases in farrowed sows by several months, demonstrating the potential of effluent monitoring as an early warning system for JEV outbreaks in piggeries. Positive effluent detections on 21/01/2025, 18/02/2025 and 11/03/2025 align with the mating and early gestation periods of the 3 suspect litters born with significant numbers of stillborn and mummified piglets on 17/05/2025. Based on mating records the estimated JEV infection exposure period for those sows was between 23/02/2025 and 30/03/2025. The 1 confirmed JEV positive litter born on 01/06/2025, had an estimated exposure period from 13/03/2025 to 17/04/2025 as per Table 2. This suggests effluent surveillance can detect JEV infections 2–4 months before clinical or reproductive impacts manifest, as evidenced by the confirmed case on 03/06/2025. The early detections in effluent may represent asymptomatic JEV activity in the herd, rather than the exposure events that led to the affected litters born in May–June. These findings support integrating RT-PCR-based effluent monitoring into routine piggery surveillance (i.e., seasonal climate and mosquito surveillance) to enable early intervention with preventative measures, such as mosquito control or herd vaccination, to be initiated to mitigate JEV risks.

The effluent detection on 11/03/2025 aligns closely with the estimated exposure window for the suspect litters born in May (estimated exposure 23/02/2025 to 30/03/2025). This same detection also falls just outside, but near, the estimated exposure period for the confirmed JEV affected litter born on 01/06/2025 (with an estimated exposure between 13/03/2025 and 17/04/2025). Nonetheless, effluent RT-PCR positives should be interpreted as early-warning signals rather than definitive evidence of sow-level exposure. Confirmation via repeat detection, increasing viral load, and triangulation with oral-fluid serology/RT-PCR, gestation timing, and vector/activity data is essential to distinguish background circulation and subclinical shedding from epidemiologically meaningful exposure events.

The intermittent nature of the detections in effluent, with negative results on 17/12/2024, 04/02/2025, 11/02/2025 and 25/02/2025, may reflect genuine non-infection of the sampled pig population, or low JEV RNA concentrations in the effluent samples (i.e., below the RT-PCR assay’s limit of detection). Additionally, the use of weekly grab sampling, which provides only a snapshot of effluent composition, may have contributed to non-detections due to diurnal variations in viral shedding or uneven JEV distribution in effluent. It is also possible that JEV was not present in the effluent sample at the time of sampling. However, effluent surveillance ceased after 11/03/2025 due to limited resources and the absence of apparent clinical cases at the time, a reactive study design that restricted data collection during the full exposure period for the confirmed June litter. This limitation highlights that the study’s design was constrained by practical considerations, potentially reducing its ability to capture the full temporal dynamics of JEV transmission and affecting the generalizability of findings to settings with continuous monitoring. As a proof-of-concept, these results underscore effluent monitoring’s potential as an early warning tool, but future studies should prioritize sustained sampling, potentially using passive samplers (Bivins et al. 2022).

RT-PCR testing of piggery effluent detected JEV on multiple occasions, while corresponding oral fluid samples from the herd returned negative results. This discrepancy suggests that oral fluid sampling may have involved only a small subset of pigs that was not reflective of the entire herd, potentially leading to undetected infections. Alternatively, if sampling involved a small proportion of pigs that were infected and secreting JEV, with the majority of contributing pigs uninfected, the aggregate oral fluid sample collected may have been diluted to levels below detectable levels. In contrast, effluent represents a composite sample from the entire herd, increasing the likelihood of detecting viral shedding. The positive effluent results on 14/01/2025, 21/01/2025, 18/02/2025 and 11/03/2025, despite negative oral fluid samples by ELISA and RT-PCR during this sampling period, indicates an enhanced sensitivity of effluent testing. Effluent surveillance thus offers a non-invasive, herd-level surveillance tool that can complement traditional veterinary clinical surveillance and testing and improve outbreak preparedness.

## CONCLUSIONS

- Effluent monitoring using RT-PCR detected JEV RNA as early as 14/01/2025, providing a 2–4-month lead time before confirmed cases on 03/06/2025, affirming its potential as an early warning system.
- The 11/03/2025 positive effluent sample aligned with the estimated JEV infection exposure period of the suspect May litters, demonstrating its ability to signal infections before reproductive impacts become evident.
- Effluent surveillance offers a non-invasive, herd-level surveillance alternative to oral fluid testing, enhancing proactive disease management in piggeries during high-risk periods.
- These findings support the integration of effluent monitoring into routine JEV surveillance to mitigate risks in Australia’s enzootic regions. While effluent sampling appears more sensitive, oral fluid testing still has value as a complementary tool. Future research should refine detection thresholds and define optimal deployment strategies.

## ACKNOWLEDGEMENTS

We thank the farm owner involved for providing samples.

